# Embryonic and larval development of the neotropical harlequin toad, *Atelopus cruciger* (Lichtenstein and Martens, 1856) (Anura: Bufonidae)

**DOI:** 10.1101/2025.09.15.675891

**Authors:** Ingrid Márquez, Celsa Señaris, Onil Ballestas, Katiuska González, Chris Buttermore, Javier Mesa, Federico Pantin, Tuenade Hernández, Margarita Lampo

## Abstract

Harlequin toads (*Atelopus*) are one of the amphibian groups in greatest need of urgent conservation measures, *Atelopus cruciger* is one of the few species with known stable populations, yet basic natural history, including the embryonic and larval stages, remains largely unknown. We describe for the first time the development of its tadpole, from egg to tail resorption. The tadpole of *Atelopus cruciger* is gastromyzophorous with a general morphology resembling that for other congeneric species but with a distinct W-shaped whitish band that spans transversely across the midline of its dorsum. Development from egg to tail resorption takes between 110–147 days. In the wild, tadpoles live camouflaged beneath and alongside rocks in shallow sections of fast-flowing streams, with clear soft waters.

## Introduction

Harlequin toads (*Atelopus* Duméril & Bibron, 1841) are one of the most threatened genera of amphibians in the world, with 93% of the assessed species listed as being at risk of extinction (Re:Wild et al., 2023, Lötters et al., 2023). Of the 99 described *Atelopus* species, three are considered extinct, and 67 are Critically Endangered, with more than half suspected to be extinct in the wild (IUCN, 2025). While most harlequin toads are in urgent need of conservation actions to avoid extinctions (Valencia & da Fonte, 2022; Lötters et al., 2023), simple natural history information such as the embryonic and larval stages of most species remain unknown. The relict status of most extant populations, the scarcity of information on the distribution and microhabitat preferences within streams have resulted in around 30% of *Atelopus* tadpoles being described (Marcillo-Lara et al., 2020; Pérez-Gonzalez et al., 2020; Lötters et al., 2022). Of nine described species known in Venezuela, five have morphological descriptions of tadpoles based on only six or less specimens (Mebs, 1980; Mijares-Urrutia & La Marca, 2005), but none have a complete description of their development.

The Rancho Grande Harlequin Toad *Atelopus cruciger* (Lichtenstein and Martens, 1856) is one of two harlequin toad species that has been rediscovered in the last 15 years in Venezuela (Rodríguez-Contreras et al., 2008; Barrio-Amorós & Torres, 2023). It disappeared in the late 1980s presumably due to chytridiomycosis, a disease caused by the fungus *Batrachochytrium dendrobatidis* (*Bd*) (Bonaccorso et al., 2003). However, two populations were discovered in 2003 after extensive explorations of its former habitats. *Atelopus crucige*r is currently listed as Critically Endangered and recommended for *ex-situ* rescue by the IUCN (Lampo et al., 2022). These relict populations have maintained a few hundred mature individuals for almost one decade (Lampo et al. 2017). Despite many hours searching streams near to sections of the river where post-metamorphic individuals are abundant, no eggs masses or tadpoles have been observed (Señaris et al., 2023). Mebs (1980) described the tadpole of *A. cruciger* from a batch of eggs spawned and reared in captivity. However, the description of later stages is lacking because all tadpoles died before development was completed. Knowledge on tadpole development times and microhabitat preferences is valuable for understanding their vulnerability to habitat loss, climate change, diseases and for designing strategies for establishing *ex-situ* back up colonies and reintroducing captive bred individuals.

Here, we provide the first complete description of the development, from egg to tail resorption, of *Atelopus cruciger* based on cohorts raised in captivity. Additionally, we present novel insights into the habitat characteristics and behavior of tadpoles in the wild.

## Material and methods

### Origin of specimens

10 adult males and 10 adult females were wild–caught between 2020 and 2022 in Cuyagua river (Estado Aragua, Venezuela) to establish a breeding colony at the Centro de Reproducción e Investigación para Arlequines (CRIA) in Caracas (**Figure 1**). This effort was part of an *ex-situ* conservation initiative aimed at securing a backup colony and producing captive bred toads to reintroduce in their historic territories. Upon arrival at CRIA, all specimens were placed in quarantine, where they were photographed, measured and weighed. To prevent contamination with *Bd* zoospores, all individuals underwent a 10–day Itraconazole treatment (Brannelly et al., 2012) and were kept in quarantine for an additional 20 days after treatment.

**Figure 1.**
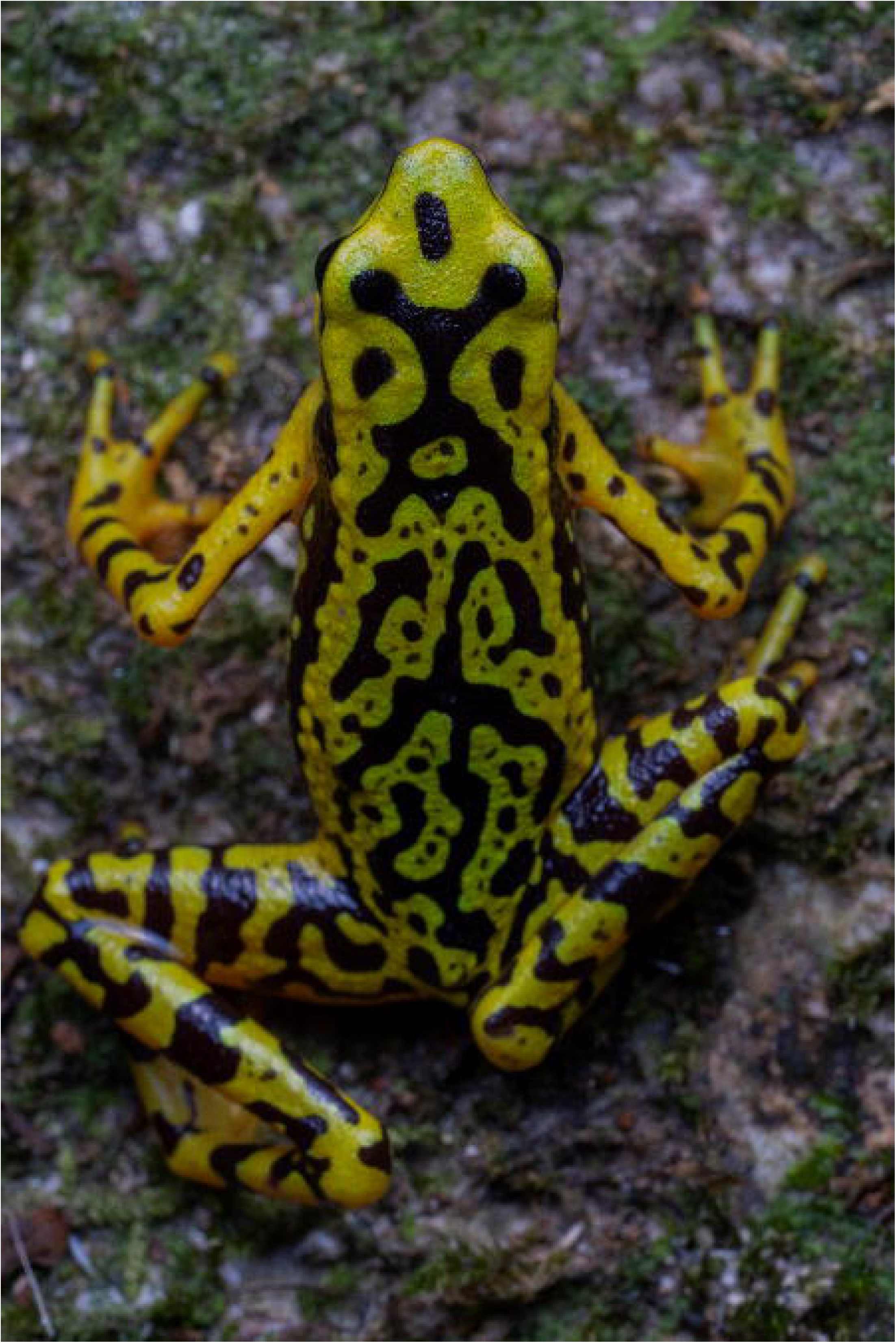
Adult female of Atelopus cruciger. Photo: Jaime Culebras

Adults were housed in 5-individual groups in 10-gallon glass tanks maintained at a temperature range of 19—22.5 °C. Their diet consisted of crickets (*Acheta domesticus*), fruit flies (*Drosophila hydei*), and fly larvae, all dusted with a supplement mixture containing calcium with vitamin D_3_, vitamin A, and a micro-fine multivitamin. Amplectant pairs were transferred to 20-gallon breeding tanks filled halfway with water and equipped with rocks of various sizes, arranged to create crevices that served as potential oviposition sites. Amplectant pairs remained in the breeding tanks for 5–12 days prior to egg deposition. After oviposition, adults were returned to their individual maintenance tanks. The first clutches were produced 7—10 months after the adults arrived at the laboratory. We obtained eggs from six clutches from four different parents. To raise tadpoles, we used reverse osmosis water reconstituted with magnesium sulfate (MgSO_4_), potassium bicarbonate (KHCO_3_), sodium bicarbonate (NaHCO_3_), and calcium chloride (CaCl_2_). Tadpoles were fed a paste with Soilent Green™ and dandelion leaves.

Additionally, during the 2020-2023 breeding seasons, we searched for and collected tadpoles along the rivers where the adult specimens were originally collected (**Figure 2**). We examined the underneath and sides of the rocks in fast-flowing waters, and backwater areas where litter accumulates along the riverbanks. A total sampling effort of 72 man-hours, tadpoles were caught by sweep-netting (21 x 15 cm). Water temperature and pH were measured at the sampling sites using a waterproof pHep+ pocket pH tester with a resolution of 0.01 pH and a precision of ±0.10. Calcium and magnesium hardness were also measured using a portable multiparameter photometer. The tadpoles collected were incorporated in the *ex-situ* program and raised to tail resorption. The tadpoles were transported to CRIA in plastic containers filled with water at 20°C, equipped with air diffusers. At CRIA, the tadpoles were kept in 20-gallon containers and fed the same diet as captive bred tadpoles. To obtain ontogenetic development data, we made daily observations on six eggs clutches laid in captivity on January 1 2023, February 1 2023, and July 19 2023, October 8 2023, December 10 2023 and February 6 2024. The specimens were deposited in the herpetology collection of Museo de Historia Natural La Salle (MHNLS) under the numbers MHNLS 22872 – MHNLS 22887.

**Figure 2.**
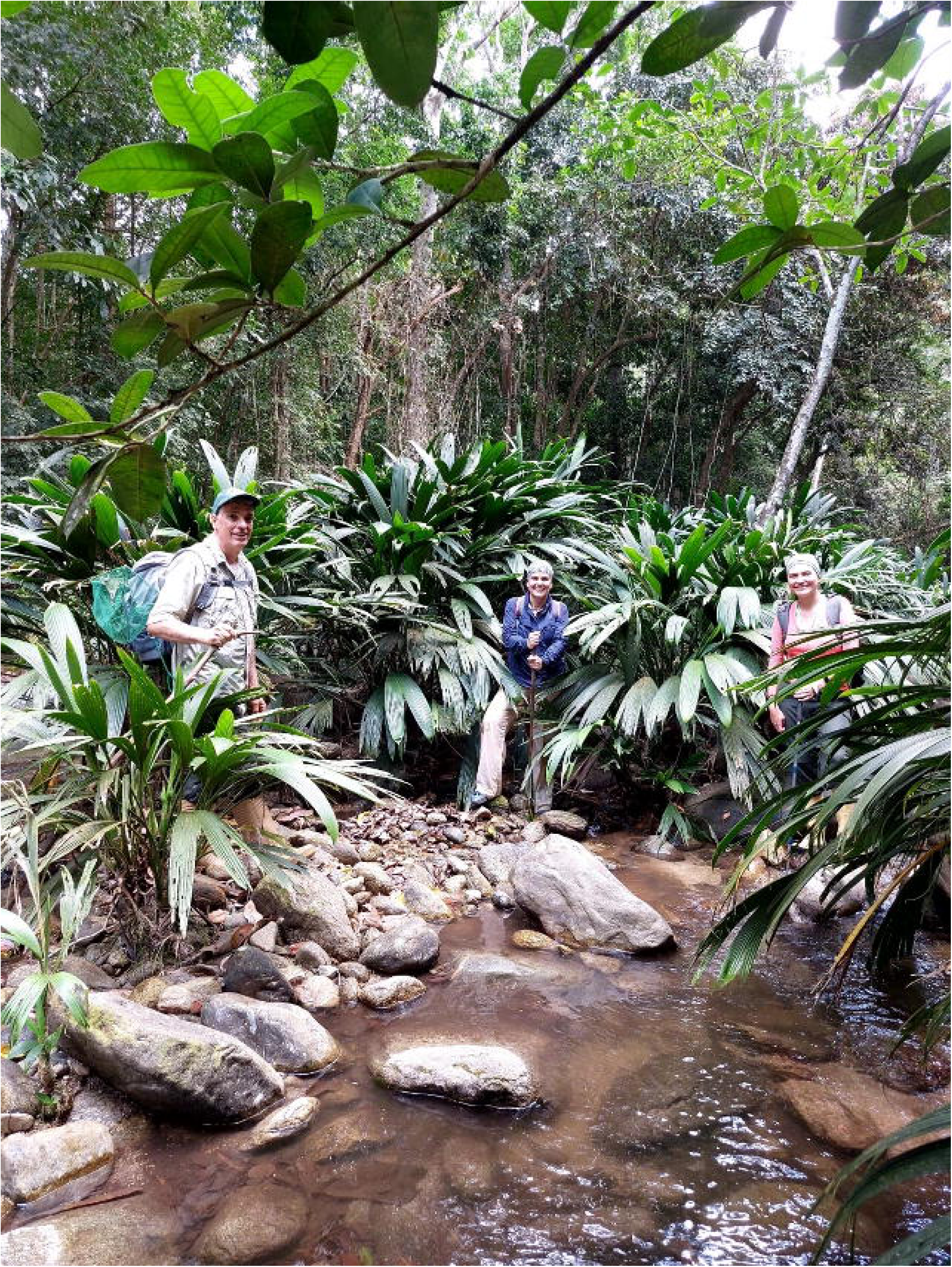
Stream section where Atelopus cruciger tadpoles have been collected by the authors. Photo: Tuenade Hernandez.

### Tadpole preservation and description

Individuals that died naturally at CRIA during its development were preserved in formalin 10% and used in this study. Additionally, we euthanized 70 individuals in stages from 17—46, to complete the development series. This procedure was carried out by immersing the tadpoles in a solution of water and 2% lidocaine until they fell asleep and died. Tadpole description follows the terminology and definitions of Kaplan (1997) and Altig (2007), the developmental stages by Gosner (1960) and the four major developmental categories by McDiarmid & Altig (1999): embryo (Gosner stages 1–19), hatchling (stages 20–25), larva (stages 26–41), and metamorph (stages 42–46). We used the following measurements: total length (TL), body length (BL), tail length (TAL), internarial distance (IND), interorbital distance (IOD), maximum tail height (MTH), tail muscle height at base (TMH), tail muscle width at base (TMW), oral disc width (ODW), and maximum width of the abdominal sucker (AS). Measurements were taken to the nearest 0.1 mm using Leica M125 Stereo Microscope with 10X/23 eyepiece and 0.8X lp/mm resolution. To document the development, growth, and coloration at different development stages, we took sequential digital photographs using a Canon EOS 100 equipped with an RF 100mm F2.8 macro lens.

## Results

### Habitat description

Three tadpoles (Stages 26—29) were found beneath and on the side surfaces of rocks, in sections of the river with fast flowing waters at depths between 30 and 60 cm. In the middle of the dry season, during April, the river measured approximately 4 meters in width and received direct sunlight for only about two hours around midday. Its banks are covered by a dense gallery forest. Water was clear, soft (calcium hardness<<1 and magnesium hardness=85.33 mg/L; SD=5.03) and neutral (pH=6.96; SD=0.05), with temperatures of 21.7°C; SD=0.52. Due to their cryptic coloration, which closely matches the surrounding rock surfaces, tadpoles were not readily visible. Therefore, we used aquarium nets to scrape the rocks and capture them.

### Embryo and hatchling description

The size of egg clutches varied, with an estimated range of 300—800 eggs. The eggs are unpigmented, approximately 1.2—1.5 mm in diameter, covered each one by a transparent jelly capsule and arranged in a single string joined by a thin membrane (**Figure 3A**). The egg string is indented, with capsule clearly differentiated. Eggs were deposited under the stones. We did not observe empty capsules but estimated that approximately 15—25% of eggs in the first clutch and 5—10% in the second clutch did not develop and were infected by mold.

**Figure 3.**
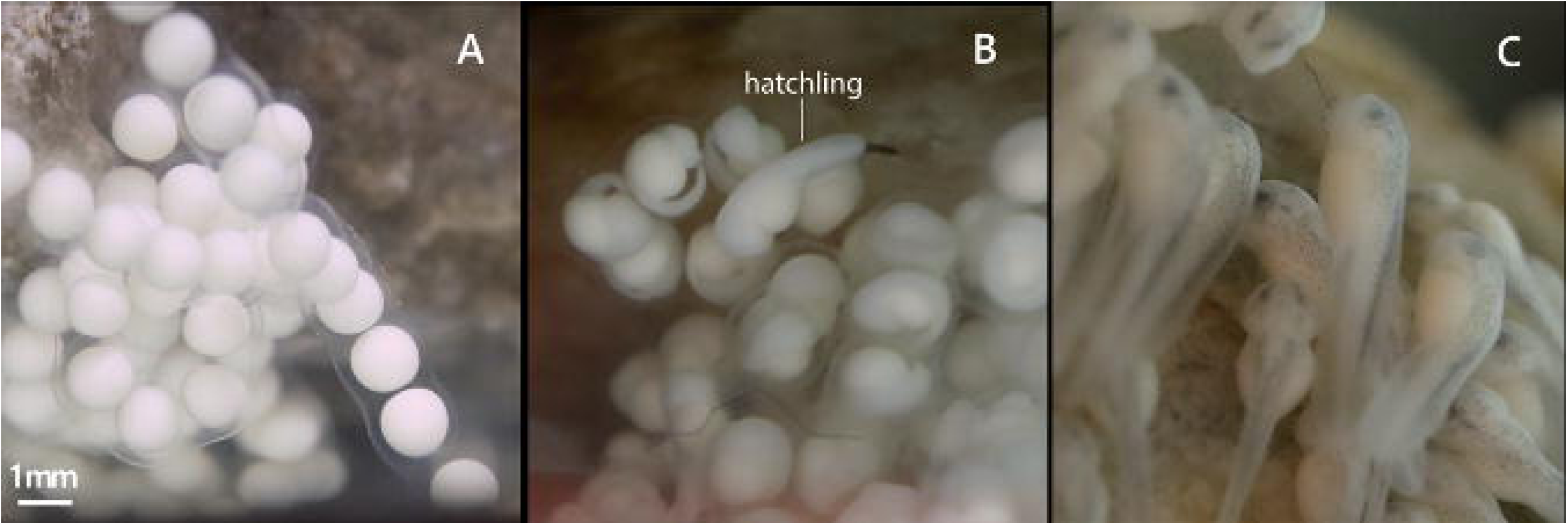
Embryonic and hatchling development of Atelopus cruciger. (A) String of white-colored eggs surrounded by a transparent gelatinous capsule and a thin membrane (day 1), (B) embryos (stages 18-19) and hatchling (day 7), and (C) hatchlings with grey pigmentation in dorsal surfaces, prospective eye regions and abdominal suctorial discs (stages 20-21) (day 10). Photos: Margarita Lampo.

Two days after the beginning of our observations, the embryos reached stage 12, and shortly after neural plate became visible on the dorsal area. Subsequently it became flat, the embryos became slightly elongated and neural folds were evident (stage 14). On day four, the neural tube was visible (stage 15-16) and head, gill plates, and the tail buds were distinguishable in most embryos by day five (stage 17). First tadpoles hatched on day six (stage 18) and muscular response was observed on that day (**Figure 3B**). A prospective eye region, a discernible abdominal suctorial disc and minute grey spots were visible on the dorsal surface on day nine (stages 19-21) (**Figure 3C**). Measurements are shown in Table 1.

**Table 1.** Measurements (mm) of tadpoles of Atelopus cruciger (mean ± SE; range in parentheses) in different development stages sensu Gosner (1960). Total length (TL), body length (BL), tail length (TAL), internarial distance (IND), interorbital distance (IOD), maximum tail height (MTH), tail muscle height at base (TMH), and tail muscle width at base (TMW), oral disc width (ODW), and maximum width of the abdominal sucker (AS).

| Stage | N | TL | BL | TAL | IND | IOD | MTH | TMH | TMW | ODW | AS |
| --- | --- | --- | --- | --- | --- | --- | --- | --- | --- | --- | --- |
| 17 | 6 | 3.36±0.21<br>(3.08–3.64) | 1.97±0.21<br>(1.60–2.20) | 1.39±0.32<br>(0.88–1.80) | | | | | | | |
| 18 | 1 | 3.60 | 1.92 | 1.68 |  |  | 0.60 | 0.40 | 0.68 |  |  |
| 20 | 1 | 5.00 | 2.40 | 2.60 |  | 0.72 | 0.60 | 0.48 | 0.56 |  |  |
| 23 | 16 | 5.39±0.92<br>(3.88–6.64) | 2.28±0.33<br>(1.68–2.96) | 3.11±0.75<br>(1.80–4.24) | 0.31±0.07<br>(0.20–0.44) | 0.54±0.13<br>(0.36–0.84) | 0.74±0.17<br>(0.44–0.96) | 0.67±0.17<br>(0.32–0.88) | 0.58±0.11<br>(0.32–0.76) | | |
| 24 | 11 | 5.94±0.72<br>(4.56–6.72) | 2.38±0.28<br>(1.88–2.60) | 3.57±0.51<br>(2.68–4.24) | 0.32±0.10<br>(0.20–0.36) | 0.55±0.13<br>(0.40–0.84) | 0.95±0.18<br>(0.64–1.20) | 0.65±0.19<br>(0.28–0.84) | 0.58±0.15<br>(0.44–0.96) | | |
| 25 | 14 | 6.19±0.57<br>(5.36–7.20) | 2.19±0.38<br>(1.60–3.12) | 4.00±0.49<br>(3.04–4.60) | 0.33±0.07<br>(0.24–0.48) | 0.47±0.15<br>(0.40–0.76) | 0.84±0.13<br>(0.68–1.12) | 0.68±0.14<br>(0.36–0.80) | 0.58±0.10<br>(0.44–0.80) | | |
| 27 | 1 | 11.44 | 4.88 | 6.56 | 0.44 | 0.68 | 1.28 | 1.0 | 1.4 |  |  |
| 28 | 1 | 10.36 | 3.96 | 6.40 | 0.50 | 0.77 | 1.80 | 1.48 | 1.09 |  |  |
| 29 | 2 | 13.24±0.65<br>(12.78–13.70) | 5.72±0.46<br>(5.39–6.04) | 7.53±0.19<br>(7.39–7.66) | 0.80±0.12<br>(0.71–0.88) | 0.96±0.18<br>(0.83–1.08) | 2.14±0.23<br>(1.97–2.30) | 1.58±0.48<br>(1.24–1.92) | 1.51±0.25<br>(1.33–1.68) | 1.05 ± 0.2<br>(0.89–1.20) | 4.02 ± 0.1<br>(3.95–4.09) |
| 31 | 1 | 14.57 | 6.00 | 8.57 | 0.65 | 1.43 | 2.67 | 1.82 | 1.73 | 1.55 | 4.73 |
| 36 | 5 | 15.03±1.3<br>(13.12–14.66) | 5.90±0.22<br>(5.74–6.05) | 8.00±1.31<br>(7.07–8.92) | 1.13±0.06<br>(1.09–1.17) | 1.59±0.74<br>(1.07–2.11) | 2.49±0.06<br>(2.45–2.53) | 1.99±0.18<br>(1.86–2.11) | 1.81±0.45<br>(1.49–2.12) | 1.48 ± 0.3<br>(0.91–1.79) | 4.39 ± 0.7<br>(3.49–4.89) |
| 38-39 | 1 | 15.55 | 7.31 | 8.24 | 1.15 | 2.02 | 2.41 | 1.85 | 1.54 | 1.56 | 5.30 |
| 40-41 | 3 | 17.00 ± 0.6<br>(16.33–17.59) | 7.35 ± 0.3<br>(7.14–7.68) | 9.65 ± 0.5<br>(9.10–9.93) | 1.13 ± 0.01<br>(1.12–1.14) | 2.32 ± 0.3<br>(2.01–2.60) | 2.85 ± 0.2<br>(2.68–3.00) | 2.15 ± 0.1<br>(2.06–2.23) | 2.09 ± 0.1<br>(1.93–2.25) | 1.75 ± 0.1<br>(1.64–1.89) | 5.29 ± 0.4<br>(4.96–5.70) |
| 42-46 | 7 | 16.26 ± 0.9<br>(15.47–17.63) | 8.10 ± 0.7<br>(7.10–9.17) | 8.16 ± 1.0<br>(6.89–9.60) | 1.28 ± 0.2<br>(1.03–1.59) | 2.51 ± 0.4<br>(1.67–2.79) | 2.38 ± 0.9<br>(2.08–2.73) | 1.85 ± 0.2<br>(1.66–2.12) | 1.70 ± 0.3<br>(1.25–2.13) | 1.22 ± 0.8<br>(0.39–2.07) | 5.44 ± 2.1<br>(4.71–6.00) |

### Larval development

The larval growth and development rates varied greatly between individuals and clutches (**Figure 4**). On day 13 post-deposition, tadpoles began to swim and adhered to rock surfaces and the tank glass with the abdominal sucker (stages 21—22). The abdominal sucker occupied one half of the abdomen, and eyes were visible. On day 18, tadpoles dispersed and fed throughout the tank; oral discs could be distinguished, and intestinal coils were visible through the integument (stage 23—24). On days 21—22, dark spots became denser and darker in dorsal surface, except in two transverse bands on both sides, where pigment a was cream color. Small golden spots scattered on the dorsal surface of the tadpoles began to appear, and golden transverse bands delineating an “W-shape” pattern at the midpoint of body (stage 25) (**Figure 5A—C**). The vent tube was fully developed, and the heart was visible through the ventral skin (**Figure 5D**). Between days 13 and 22 (stages 18—25) ∼50% of the tadpoles from the first clutch and ∼10% of the tadpoles from the second clutch had died and were collected for inspection.

**Figure 4.**
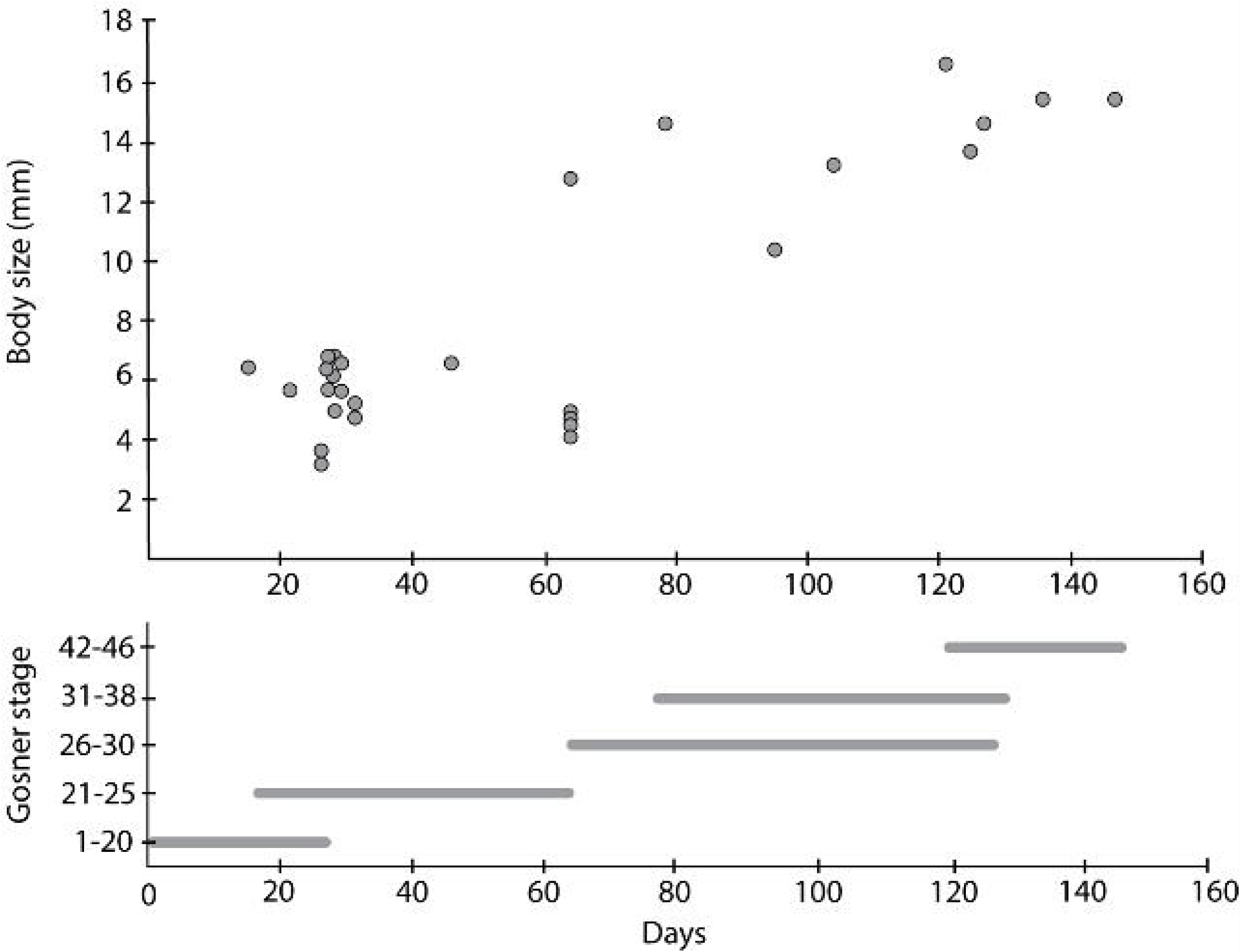
Larval growth and development of Atelopus cruciger.

**Figure 5.**
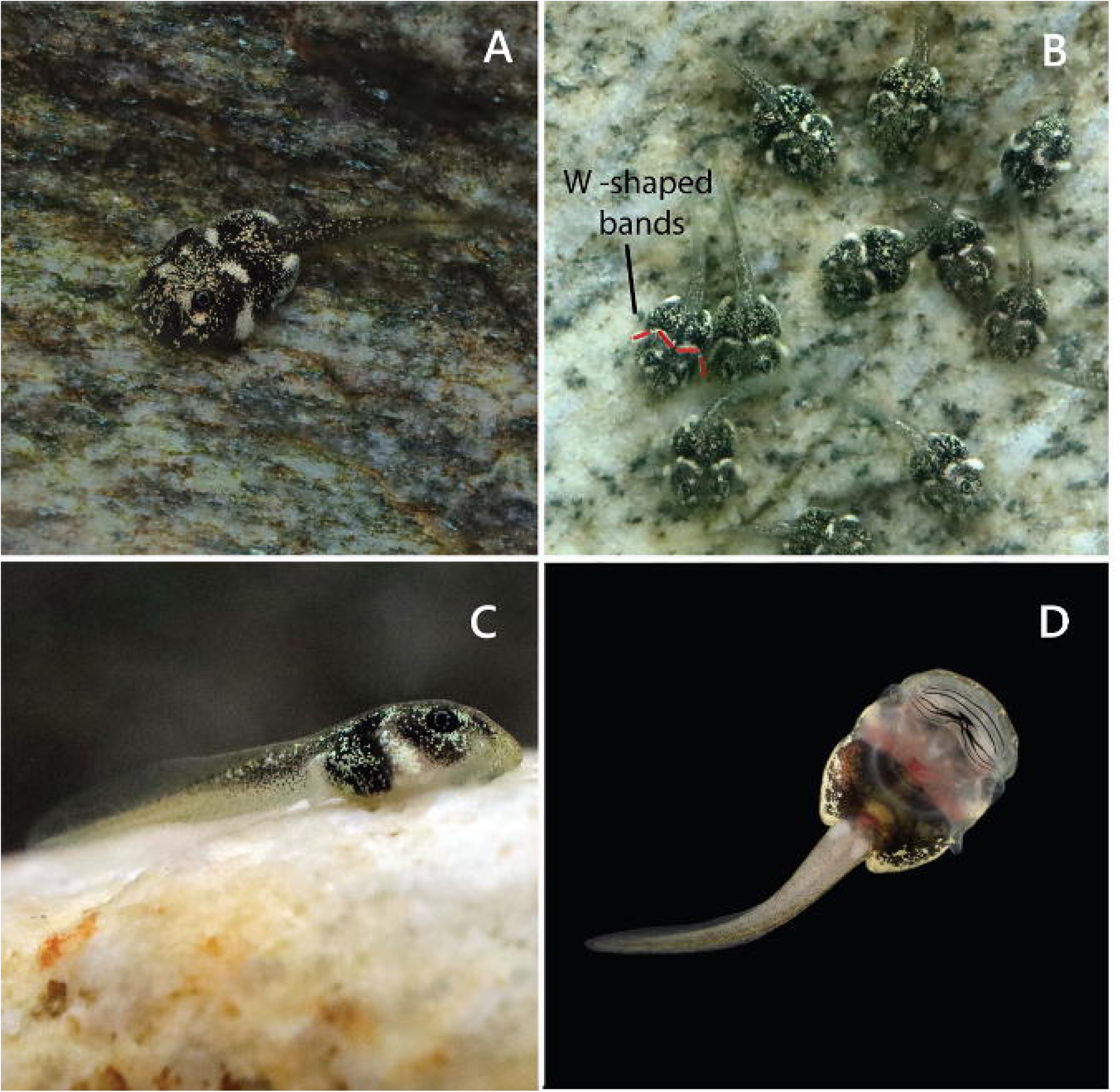
Tadpoles of Atelopus cruciger on (Stages 25—28). A) Dorsal view. B) Characteristic W-shaped whitish dorsal and lateral segmented band at midbody. C) Lateral view. D) Ventral view (stage 28). Photos: Margarita Lampo (A, B, C) and Jaime Culebras (D).

At day 55, we began to observe hindlimb buds in larger tadpoles (stage 26), and on day 78, buds were 1.5x long as the base diameter (stage 29) (**Figure 6A**) and the liver was visible on the right side of the abdomen anterior to the intestines. After day 78, some tadpoles began to show a «paddle-shaped» foot as described by Gosner (1960) was visible (stage 31) and toes 2—5 were well defined around day 100 (stage 36) (**Figure 6B**). On day 104, the forelimbs, still enclosed in the branchial chamber, were visible through the integument (**Figure 7A**), and 3—5 days later, the new mouth develops and the toadlet is ready to leave the water (**Figure 7B**).

**Figure 6.**
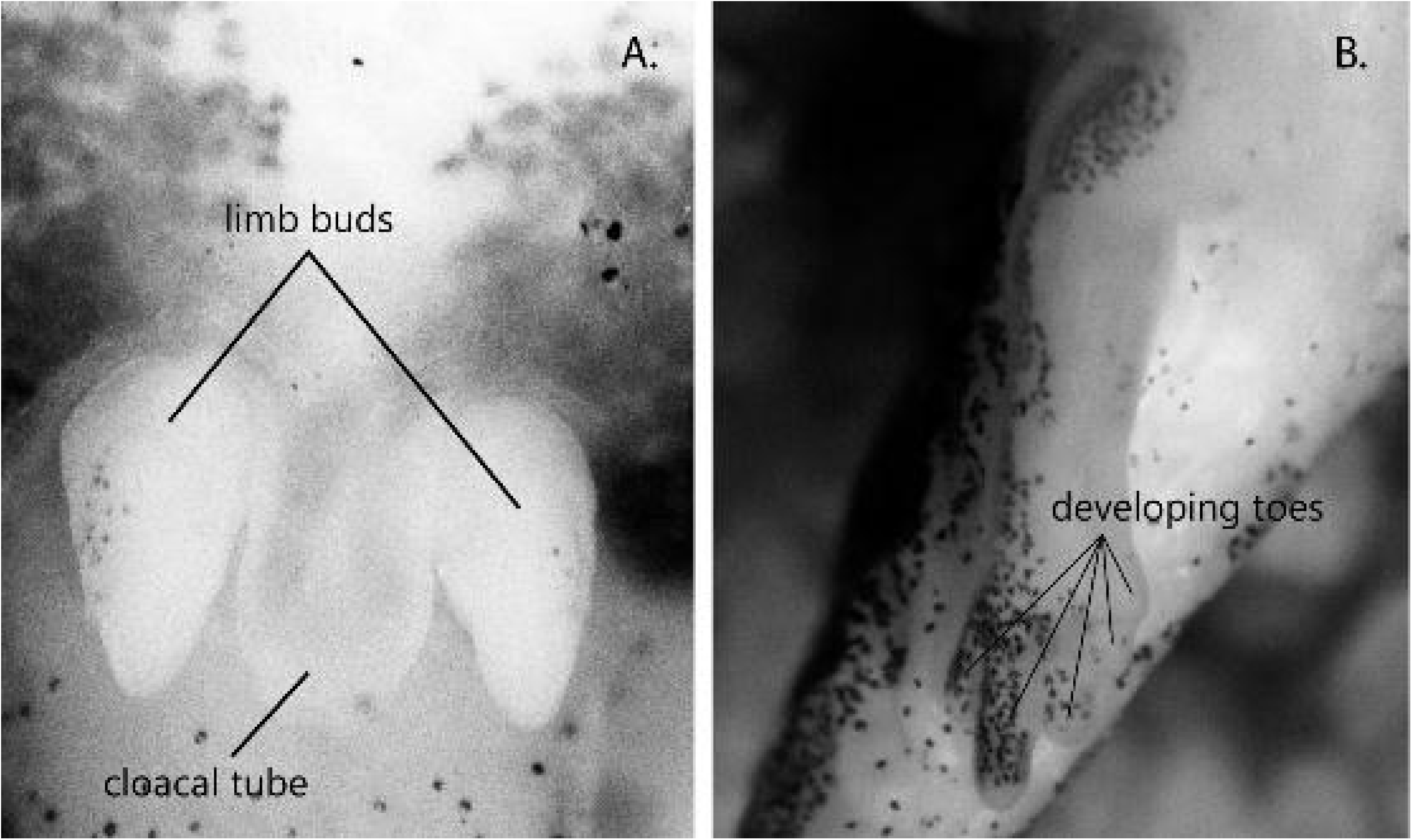
Tadpole of Atelopus cruciger A) Limb buds and cloacal tail piece at stage 29. B) Developing toes at stage 36; Toes 2—5 well defined, toe 1 incipient. Photo: Ingrid Márquez.

**Figure 7.**
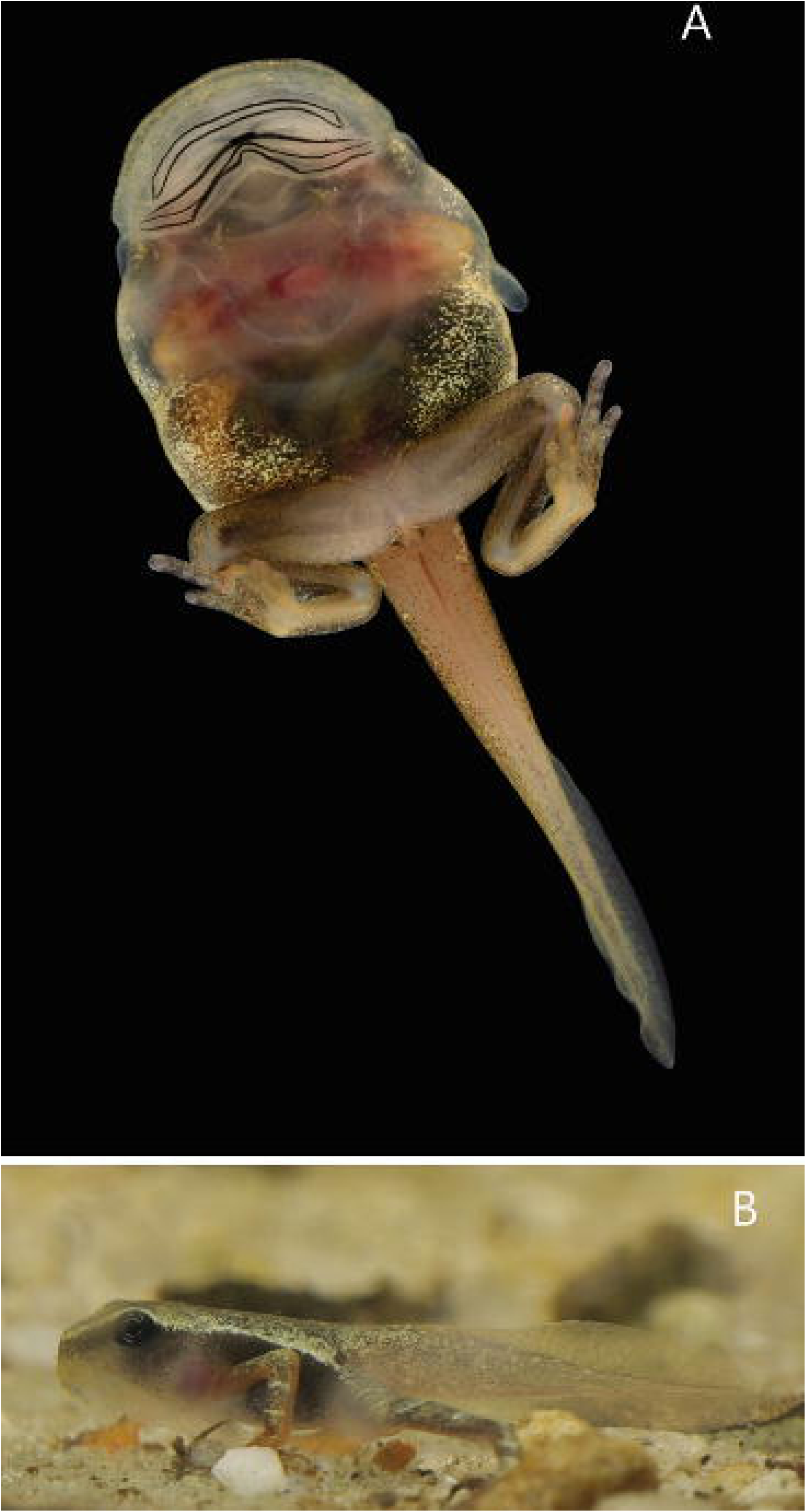
A) Ventral view of stage 40-41 larvae. Forelimbs are visible through the transparent skin and ventral tube still present. (B) Lateral view of a metamorph (Stage 43): translucent membrane of dorsal and caudal fins visible. Forelimbs complete and mouth corner anterior to eye. Photos: Jaime Culebras (A) and Margarita Lampo (B).

### Tadpole description

The following description is based on 14 individuals in stage 25. Measurements (in mm) are as follows (see also Table 1): TL= 6.19±0.57; BL= 2.19±0.38; TAL = 4 ± 0.49; IND = 0.33 ± 0.07; IOD = 0.47 ± 0.15; MTH = 0.84 ± 0.13; TMH = 0.68 ± 0.14; TMW = 0.58 ± 0.10. Body ovoid in dorsal view, broader at the midpoint -posterior to eyes, slightly depressed in lateral view, and ventrally flattened with the tail clearly separated; snout broadly rounded in dorsal view, and rounded or slightly curved from the side profile; chondrocranial elements not visible; eyes positioned dorsolaterally, the distance between them approximately two times their diameter and about 1.6 times the internarial distance; nostrils small, rounded to ovoid, with non-protrusive margin, visible in dorsal and lateral views, closer to eyes than the tip of snout. Spiracle sinistral, relatively small and cylindrical, single, unpigmented, originating at midpoint of body, not attached to body, and opening directed posterolaterally. Vent tube tubular, and narrow, medial, linked to the caudal muscle, directed posteriorly opening with a medial horizontal notch. Caudal musculature moderately weak, keeping its size in the proximal half and decreasing quickly in distal half, not reaching tip of tail. Relative tail length variable, from 1.3 to twice the body length, TAL mean 61.8% of TL; caudal musculature robust in anterior third of its length, narrowing quickly posterior to midpoint of tail, close to but not reaching tip of tail. Dorsal and ventral fins originating on the tail, notably narrower than caudal musculature in the anterior half of tail length; tail tip rounded. No lateral line or glands detectable. Mouth ventral, surrounded by well-developed labia forming complete not emarginated oral disc, as wide as the body, bordered by a row of short and rounded papillae interrupted by a large ventral gap of about 75-80% of the oral disc width (**Figure 8**). Rostral papillae slightly larger and better defined than lateral ones, submarginal papillae apparently absent. Labial tooth row formula 2/3; all teeth approximately equal in length, upper tooth rows (A1, A2) slightly smaller than lower tooth rows, occupying nearly the entire width of the oral disc. Jaw sheaths well keratinized, serrated; upper jaw sheath arch-shaped and shorter than the V□shaped lower sheath. Abdominal sucker large, extending from posterior labium to two-thirds of the body length, circular or slightly ovoid at posterior end, occupying around 70— 78% of body width; lateral and posterior edges of abdominal sucker free from body.

**Figure 8.**
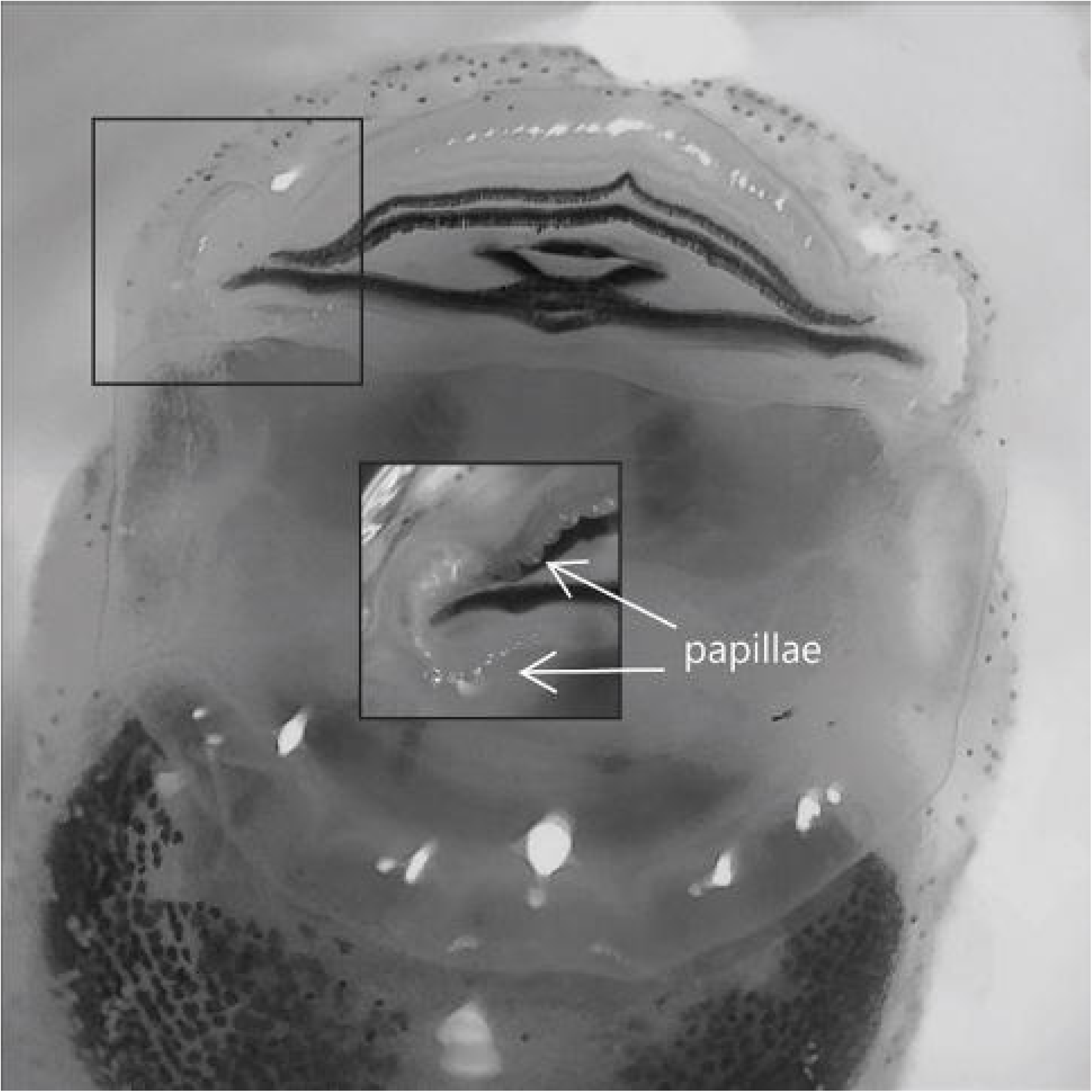
Ventral view of the oral disk of a tadpole of Atelopus cruciger (Stage 25). Posterior (lower) tooth rows (P1 –P3) are collapsed in the photo. Marginal papillae are shown in the detail square. Photo: Margarita Lampo.

### Color in life

Dorsum and flanks black with a conspicuous whitish lateral band and two dorsal blotches at midbody that form an inverted, segmented W-shape and minute creamy and golden metallic spots, most densely concentrated around the eyes and the posterolateral end of the body (**Figure 5**). Spiracle completely translucent. Tail musculature anterodorsally black with whitish and golden metallic spots, and the rest cream with scattered, minute, dark spots; dorsal and ventral fins translucent (**Figure 5C**). Belly dark, with whitish and golden minute spots on sides, anterior part unpigmented and translucent. Oral disc and abdominal sucker translucent. Jaw sheaths and labial teeth black (**Figure 5D**). Eye black, with a light golden ring around the pupil.

During ontogeny, and from stage 41, the dorsum becomes dark brown with small cream spots, and a conspicuous fine whitish line extended laterally from the tip of the snout, above the eyes, to the groins. Tail musculature becomes light brown and yellowish by stages 41-43 (**Figure 7B**). Also, a faint light irregular banding appears on a brown background in the forehands and hindlimbs, which persists until metamorphosis completed (Stage 46).

### Color in preservative

Color pattern is similar to that of living tadpoles but fades and loses the golden and iridescent tones; whitish spots and tail musculature becomes light cream. Spiracle unpigmented. The intestine is visible (dark) in the translucent cream belly. Eye black, pupil dark grey.

## Discussion

This study presents the first comprehensive embryonic and larval staging for *Atelopus cruciger*, shedding light on the largely unknown early life stages of this critically endangered species. These findings are crucial for identifying *A. cruciger* individuals in their early life stages in the wild and can provide new insights into the taxonomy and evolution of this genus (Lötters et al., 2022; dos Santos Dias & Anganoy-Criollo, 2024). Additionally, our results also present a timeline in the sequential Gosner stages —from eggs to tail resorption— and larval growth, valuable for better understanding the population dynamics of the species, its mechanisms for coexistence with *Bd* (Lampo et al., 2017; Ballestas et al., 2021), and the environmental cues that trigger reproduction in the wild.

Development time, from egg to tail resorption, in *A. cruciger* took between 110—147 days at 20—22 °C, and toadlets emerged from the water with a body size (SVL) between 7.1—9.2 mm, slightly smaller than observations in the wild of 10—12 mm (Señaris et al., 2023). To our knowledge, only three other *Atelopus* species, all from ex situ conditions, have larval development chronologies. Development time ranged from 115 to over 200 days in *Atelopus balios* Peters, 1973, 133–140 days in *Atelopus flavescens* Duméril & Bibron, 1841, later rediscovered as *Atelopus hoogmoedi nassaui* (Gawor et al., 2012; Lötters et al., 2022; Buttermore et al., 2024) and between 120 and 240 days in *Atelopus zeteki* Dunn, 1933 (Poole, 2006). This means that under experimental conditions, the larval development time —aquatic phase— in *Atelopus* exhibits a wide range, but averages between 3.5 and 4.5 months, possibly due to husbandry conditions (*e.g.* temperature, density, nutritional condition) or genetic predetermination. Egg hatching and tadpole survival varied significantly between clutches, even under similar water and nutritional conditions. It is unclear whether this variability also occurs in the wild.

The general morphology of the tadpole of *Atelopus cruciger* resembles that for other congeneric species (Marcillo-Lara et al., 2020; Lötters et al., 2022; dos Santos Dias & Anganoy-Criollo, 2024), yet it exhibits a unique combination of characteristics specific to this species. It is a gastromyzophorous type of exotrophic anuran larvae, with a large abdominal sucker and robust caudal musculature adapted for attaching to rocks and swimming in fast-flowing waters, a sinistral spiracle, and an oral disc with marginal papillae and a labial tooth row formula 2/3, this dental formula is shared in the genus *Atelopus*; the variations are due to the shape of the bill, which is concave at the top, while in *A. tricolor* it is convex. In species from the Venezuelan Andes, the bill is concave or slightly concave (Mijares-Urrutia & La Marca, 2005). However, some characteristics may be useful to distinguishing the larvae of *Atelopus* species as the color pattern of the body and tail, the presence or absence of submarginal papillae, and size at different stages (Marcillo-Lara et al., 2020), but also the relative size of the abdominal sucker, the tail length relative to total length, the relative lengths of the upper and lower jaw sheaths, and the size of the spiracle (Pérez-Gonzalez et al., 2020; Lötters et al., 2022).

The majority of the known *Atelopus* tadpoles are uniformly dark, brown or black, without contrasting marks (Lötters, 2001; Boistel *et al*., 2005; Marcillo-Lara *et al*., 2020), while other species show striking whitish or cream spots, bands, or blotches (Pérez-Gonzalez *et al*., 2020). The larva of *A. cruciger* can be identified by a distinct W-shaped whitish band that spans transversely across the midline of its dorsum. The absence of a black band on the fins and caudal musculature is also characteristic of this species, although this trait is shared with *A. carbonerensis* Rivero, 1974, *A. certus* Barbour, 1923, *A. coynei* Miyata, 1980, *A. mittermeieri* Acosta-Galvis et al., 2006, *A. nahumae* Ruiz-Carranza et al., 1994, *A. palmatus* Andersson, 1945, *A. sorianoi* La Marca, 1983, and *A. subornatus* Werner, 1899; It contrasts with A. *sonsonensis*, which has a band of dark brown chromatophores in the caudal musculature (Vélez-Rodríguez & Ruíz-Carranza, 1997). This cryptic coloration, in addition to its small size, makes this tadpole virtually invisible to the unaided eye against the rock background in the fast-running waters (Karraker *et al*., 2006).

The tadpole of *A. cruciger* has a relatively large abdominal sucker (about 65-68% of the body length), a feature shared only with *A. elegans* Boulenger, 1882, *A. nanay* Coloma, 2002, *A. subornatus* and *A. tamaense* La Marca et al., 1990, and which differs from what was found by Mijares-Urrutia and La Marca (2005) in *A. sorianoi* which has a larger abdominal sucker (about 73% of the body length). Regarding the caudal musculature, *A. cruciger* resembles *A. sorianoi* since the distal end of the tail is oval, the caudal musculature does not reach the distal end, and the caudal fins have their origin in the tail. Submarginal papillae had been reported in 40.7% species (Marcillo-Lara et al. 2020), but this character apparently varies ontogenetically and is present in advanced stages tadpoles (stage 30). In the *A. cruciger* tadpoles examined, no submarginal papillae were observed at any stage of development; however, their presence cannot be ruled out.

A recent phylogenetic study suggested that *A. cruciger* belongs to the same phylogenetic clade as the Venezuelan Andean species (Lotters et al., 2025). Although morphologically, *A. cruciger* tadpoles share features with *A. tamaense*, *A. carbonerensis*, and *A. sorianoi*, such as caudal musculature and the shape and location of the spiracle, they also share features with *A. certus* and *A. sobornatus*, which belong to the Andean-Choco-Central American clade, such as the absence of a black band on the fins and caudal musculature. Additionally, *A. cruciger*, like *A. sorianoi*, do not have the lateral body constrictions present in *A. carbonerensis*, *A. mucubajiensis* and *A. tamaense*. The Venezuelan Andean species would need to be included in phylogenetic studies to make inferences regarding phylogeny and morphology. Mijares-Urrutia and La Marca (2005) suggest that certain morphological features, such as coloration musculature, may be more related to the environment in which the tadpoles develop than to phylogeny.

Since it is difficult to collect tadpoles, it has not been possible to monitor them in the wild, which is why observations of captive-bred individuals provide knowledge about the life history of rare or endangered species. These observations give insights into evolutionary relationships with other species of the genus and are valuable for developing strategies to mitigate threats and restore decimated populations (Gawor et al., 2012; Lötters et al., 2023; Buttermore et al., 2024). For example, optimizing breeding conditions, detecting health issues, identifying most vulnerable stages and deciding optimal release strategies in reintroduction programs (Canessa et al., 2014). Although further investigations into this critically endangered species are necessary, understanding the larval development provides opportunities to build upon with regards to the species’ specific conservation needs. From the results of this work, we are prioritizing protection of the riparian habitats needed to ensure developmental success in this species which is vital to sustaining and improving the wild population. Aside from the management of the natural habitat of this species, we will begin to focus on refining our husbandry regimes so that liberation of captive bred individuals into the wild can be a viable option in the near future.

The scourge of the chytrid fungus, which is presumed to be a factor in the initial demise of *A. cruciger*, continues across the world and is a determining factor in the viability of reintroductions as a recommended conservation actions in other *Atelopus* species; most notably, *A. zeteki* which has existed in captive breeding programs for more than two decades (Poole, 2006). Assessments on the current presence or absence of *B. dendrobatidis* in the native range of *A. cruciger* are a necessity towards realizing the full potential of the *ex-situ* work we have started in this publication. While we now have a successful methodology of captive production of this species, and an understanding of the normal larval development cycle, we must gauge the ability of these captive bred individuals to survive in the wild. Understanding the fitness of captive produced *A. cruciger* released into their native ranges is a crucial step to improving the success of this conservation project and thus the survival outcomes of this species.

The high variability in survival between clutches observed in *A. cruciger* has previously been reported in captive-bred larval *Atelopus* (Gawor et al., 2012; Augustine et al., 2023), presenting an opportunity to explore ways to enhance captive production and the quality of specimens produced *ex-situ*. While this variability may represent an inherent developmental pattern in certain *Atelopus* species, it could also be associated with the conditions of the artificial streams constructed for their reproduction in captivity or other aspects of their care. It is important to deepen our understanding of the mechanisms influencing offspring survival, specifically whether they selectively favor the fittest tadpoles or operate indiscriminately, merely limiting the overall production of captive-bred individuals.

Fifteen years ago, *Atelopus cruciger* was believed to be extinct, but today it is the focus of extensive conservation efforts aimed at fully understanding its status in the wild (Rodríguez-Contreras et al., 2008), preserving its genetic pool ex-situ, and supporting wild populations with individuals bred in controlled environments. While there is still much to learn to ensure this is done responsibly, we are confident that continuing this path can lead to more direct applications of our captive assurance population and serve as a model for future amphibian conservation projects.

## Acknowledgements

This research was supported by grants by The Amphibian Ark (Grants 2022— 2023, 2023—2024). Permits were issued by Ministerio de Ecosocialismo (MINEC) (Resolución No. 008826, 22 September 2022; Resolución No. 0024, 18 March de 2022). We would like to thank Oscar Lasso-Alcalá and Mati Aristeguieta for the initial field sampling of tadpoles (2020-2021). We would like to thank Jaime Culebra for the photographic record of CRIA activities and the photos that illustrate this work. Finally, we would like to thank Jaime Nestares for his unconditional support to the *ex-situ* conservation program of *Atelopus cruciger*. Additionally, we would like to thank the reviewers of this work, Stefan Lötters and Luis Alberto Rueda-Solano, for their time, corrections, and suggestions that helped improve the manuscript.

## Notes

### Competing Interest Statement

The authors have declared no competing interest.

